# Interaction of SCoV-2 *NSP7* or *NSP8* alone with *NSP12* causes constriction of the RNA entry channel: Implications for novel RdRp inhibitor drug discovery

**DOI:** 10.1101/2023.07.26.550660

**Authors:** Deepa Singh, Tushar Kushwaha, Rajkumar Kulandaisamy, Vikas Kumar, Kamal Baswal, Saras H Tiwari, Arkadyuti Ghorai, Manoj Kumar, Saroj Kumar, Aparoy Polamarasetty, Deepak Sehgal, Madhumohan R Katika, Suresh Gadde, Marceline Côté, Sarala R Kayampeta, Mohan B Appaiahgari, Krishna K Inampudi

## Abstract

RNA-dependent RNA polymerase (RdRP) is a critical component of the RNA virus life cycle, including SCoV-2. Among the Coronavirus-encoded proteins, non-structural protein 12 (*NSP12*) exhibits polymerase activity in collaboration with one unit of *NSP7* and two units of *NSP8*, constituting the RdRp holoenzyme. While there is abundant information on SCoV-2 RdRp-mediated RNA replication, the influence of interplay among *NSP12, NSP7*, and *NSP8* on template RNA binding and primer extension activity remains relatively unexplored and poorly understood. Here, we recreated a functional RdRp holoenzyme *in vitro* using recombinant SCoV-2 *NSP12, NSP7*, and *NSP8*, and established its functional activity. Subsequently, molecular interactions among the *NSP*s in the presence of a variety of templates and their effects on polymerase activity were studied, wherein we found that *NSP12* alone exhibited notable polymerase activity that increased significantly in the presence of *NSP7* and *NSP8*. However, this activity was completely shut down, and the template RNA-primer complex was detached from *NSP12* when one of the two cofactors was present. Through computational analysis, we found that the template RNA entry channel was more constricted in the presence of one of the two cofactors, which was relatively more constricted in the presence of *NSP8* compared to that in the presence of *NSP7*. In conclusion, we report that *NSP7* and *NSP8* together synergise to enhance the activity of *NSP12*, but antagonise when present alone. Our findings have implications for novel drug development, and compounds that obstruct the binding of *NSP7* or *NSP8* to *NSP12* can have lethal effects on viral RNA replication.

## 1. INTRODUCTION

Members of the family *Coronaviridae* have become a major threat to public health in recent times, and those that are responsible for severe life-threatening diseases in humans mostly belong to the betacoronavirus genus (**1, 2**). <u>S</u>evere <u>A</u>cute <u>R</u>espiratory <u>S</u>yndrome-related <u>C</u>orona<u>V</u>irus-<u>2</u> (SARS-CoV-2 or SCoV-2) is the latest addition to this list and is the etiologic agent of the COVID-19 pandemic. Since its first occurrence in Wuhan, China, in the last quarter of 2019, the virus has spread quickly and escalated into a pandemic by early 2020. To date, the virus has accounted for >680 million cases of clinical disease and >6.8 million deaths across the world (**3, 4**). The virus is an enveloped particle measuring about 80-160 nm consisting of a single-stranded positive-sense RNA genome of about 30 kb. The viral genome has 14 open reading frames (ORFs), and together, they code for 29 proteins, including 16 non-structural proteins (*NSP 1–16*). *NSPs* play key roles in viral genome replication and transcription, as well as in escape from host antiviral defences (**5–7**).

Viral genome replication and transcription are mediated by the RNA-dependent RNA polymerase (RdRp) holoenzyme, composed of one copy each of *NSP12* and *NSP7*, and two copies of *NSP8*, which in complex with the template RNA-bound primer, form the RdRp complex (**8**). Because of its critical role in the viral life cycle as well as its high structural and sequence conservation across the *Coronaviridae* family, the RdRp complex is considered a major therapeutic target (**9, 10**). The enzymatically active *NSP12* has three important domains, including the C-terminal RdRp domain, the structure of which resembles that of the right hand, comprising fingers, palm, and thumb subdomains. The thumb subdomain binds to *NSP7* and *NSP8*-b, whereas the finger subdomain interacts with *NSP8*-a (**11–13**). The palm region is highly conserved, comprising polymerase motifs A-G that constitute the active site of the RdRp and are arranged strategically to provide four positively charged solvent-accessible paths: the template entry, the primer entry, the NTP entry and the nascent strand exit paths, within its central cavity to facilitate template-dependent RNA synthesis (**5, 14, 15**). The crystal structure of the SCoV-2 *NSP7-NSP8* complex revealed a 2:2 heterotetrameric form mediated by two distinct oligomeric interfaces. Interface I facilitates heterodimeric *NSP7-NSP8* assembly, while Interface II mediates the heterotetrameric interaction between the two *NSP7-NSP8* dimers (**16**). Also, *NSP8* assumes different conformations depending on its location within the RdRp complex, highlighting its remarkable structural flexibility (**11, 13, 17**). Together, these structural data suggest that the *NSP7/8* complex acts as a platform for efficient assembly of the RdRp complex.

During coronavirus replication, the RdRp holoenzyme carries out efficient RNA polymerase activity. It is reported that *NSP7* and *NSP8* act as cofactors and enhance the functionality of *NSP12* by facilitating interaction between *NSP12* and the template-primer RNA (**18**). However, studies on the SCoV-1 *NSP12* have contradicting data on its RNA catalytic activity in the absence of the *NSP7* and *NSP8*. Accordingly, while an early study reported polymerase activity for SCoV-1 *NSP12* in the complete absence of *NSP7* and/or *NSP8* (**19**), a later study, based on the biochemical data, reported an indispensable role for the two cofactors in the RNA polymerase activity of SCoV-1 *NSP12* (**20**). Contrary to this, studies on SCoV-2 have conclusively demonstrated that both the cofactors are critical for the polymerase activity of *NSP12* **(13, 21–23)** except for the study by Biswal et.al., where the authors, through biochemical studies, reported a weak polymerase activity for *NSP12, NSP12/7* and *NSP12/8* and concluded that formation of the RdRp holoenzyme is essential for efficient RNA replication (**16**). The ambiguity and differences in functionality between *NSP12*s of SCoV-1 and SCoV-2 are surprising, considering the 96% sequence identity between the two. Due to the critical role of the *NSP7* and *NSP8*, a comprehensive understanding of the architecture and dynamics of the *NSP7-NSP8* interaction with *NSP12* is essential to understand their influence on the polymerase activity and to develop effective antiviral therapies targeting the RdRp complex. Further, it is also important to understand the template cooperativity and the influence of different templates on the polymerase activity of the *NSP12*.

In this study, we aimed to investigate the individual contributions of *NSP12, NSP7, and NSP8* to RNA polymerase activity and the molecular mechanisms underlying the formation of the RdRp complex. First, when recombinant *NSP12, NSP7* and *NSP8* proteins were incubated with a variety of template-primer duplexes in different combinations and permutations, though relatively less efficient compared to the RdRp holoenzyme, *NSP12* alone was able to efficiently carry out RNA polymerase activity in a primer-dependent manner. Furthermore, contrary to some recent reports, the RdRp complex catalysed primer extension from the DNA template-DNA primer complex using dNTPs, but failed to do so using rNTPs (**24**). However, *NSP12* alone failed to catalyse the DNA template/DNA primer complex in the presence of rNTPs or dNTPs. The RNA-protein binding assays suggested direct binding of *NSP12* with the RNA template-primer duplex in the complete absence of the *NSP7/8* complex, which is consistent with previous reports (**23**). Interestingly, the *NSP12*-RNA duplex interaction was abrogated in the presence of *NSP7* or *NSP8* alone and/or the bound RNA duplex dissociated from the preformed *NSP12*-RNA complex when one of the two cofactors was added, suggesting that the *NSP* cofactors, in isolation, negatively regulate RNA polymerase activity. So, to understand the effect of these interactions on the structure and function of *NSP12*, we carried out Molecular Dynamic simulations analysis, wherein we found that the template entry path in *NSP12* was highly constricted in the presence of *NSP7* or *NSP8* alone compared to that in the presence of both the cofactors or in their complete absence. These findings have future implications in the discovery of novel RdRp-targeted antiviral drugs.

## 2. MATERIALS AND METHODS

### 2.1. Bacterial expression of recombinant *NSP12, NSP7* and *NSP8*

The cDNA coding for the *NSP12* of SCoV-2 was codon-optimised for bacterial expression and custom-synthesised with a C-terminal 10X-His tag in the pET22b vector. Similarly, cDNAs coding for *NSP8* and *NSP7* proteins were codon-optimised and custom-synthesised with an N-terminal 6X-His tag in the pET28a vector (**Figure S1, GenScript, USA**). The *NSP12, NSP8* and *NSP7* plasmid constructs were used to transform *E. coli* BL21 (DE3), selected clones for each construct were grown at 37ºC in LB broth supplemented with appropriate antibiotics and induced with 1 mM of isopropyl-D-1-thiogalactopyranoside (IPTG) when the OD_600_ was around 0.6. *NSP12, NSP8* and *NSP7* cultures were induced for 20hrs, 16hrs and 18hrs, respectively, at 16ºC. The induced cell pellet was harvested by centrifugation, resuspended in lysis buffer (20 mM Tris-HCl, pH 8.0, 150 mM NaCl, 4 mM MgCl_2_, 10% Glycerol), sonicated and then centrifuged to obtain the clarified supernatant. The supernatant was passed through the 0.2-micron filters and then through a pre-equilibrated His-Trap FF column. The recombinant proteins were eluted with the lysis buffer containing 300 mM Imidazole, which were further purified by size-exclusion chromatography using a HiLoad 26/600 Superdex 200 pg column (Cytiva, USA). The purified fractions were concentrated, and the purity was analysed on 12% SDS-PAGE followed by confirmation through western blot analysis using anti-His antibody and stored at - 80°C for further use.

### 2.2. Design and synthesis of nucleic acid templates

To study the template cooperativity and the influence of different types of templates on the RdRp activity, five different RNA templates and a DNA template were designed, and the same are listed in **Table 1** (**13, 19**). The 5’-FAM labelled (indicated by an asterisk, *) RNA, DNA primers and the loop-forming RNA template were commercially procured (IDT, USA). The four linear RNA templates, engineered to have a T7 promoter upstream of the template-coding sequence, were synthesised by *in vitro* transcription using the RiboMAX™ system (Promega, Cat # P1300) as per the manufacturer’s recommendations. The RNA templates were further purified using C4 RP columns (Cat # 214TP104, Vydac 214TP 10μm C4, Hichrom), the same was confirmed on 7M Urea polyacrylamide gels (14% Urea-PAGE) and quantified spectrophotometrically. To generate the template-primer complexes, 500 pmoles each of the template and the 5’-FAM-labelled primer were added to 100 μL reaction mix containing 1X annealing buffer (**AB**; 10 mM Tris-HCl, pH 8.0, 25 mM NaCl, and 2.5 mM EDTA), heated to 95°C for 7 min and gradually cooled to room temperature.

**Table 1:**
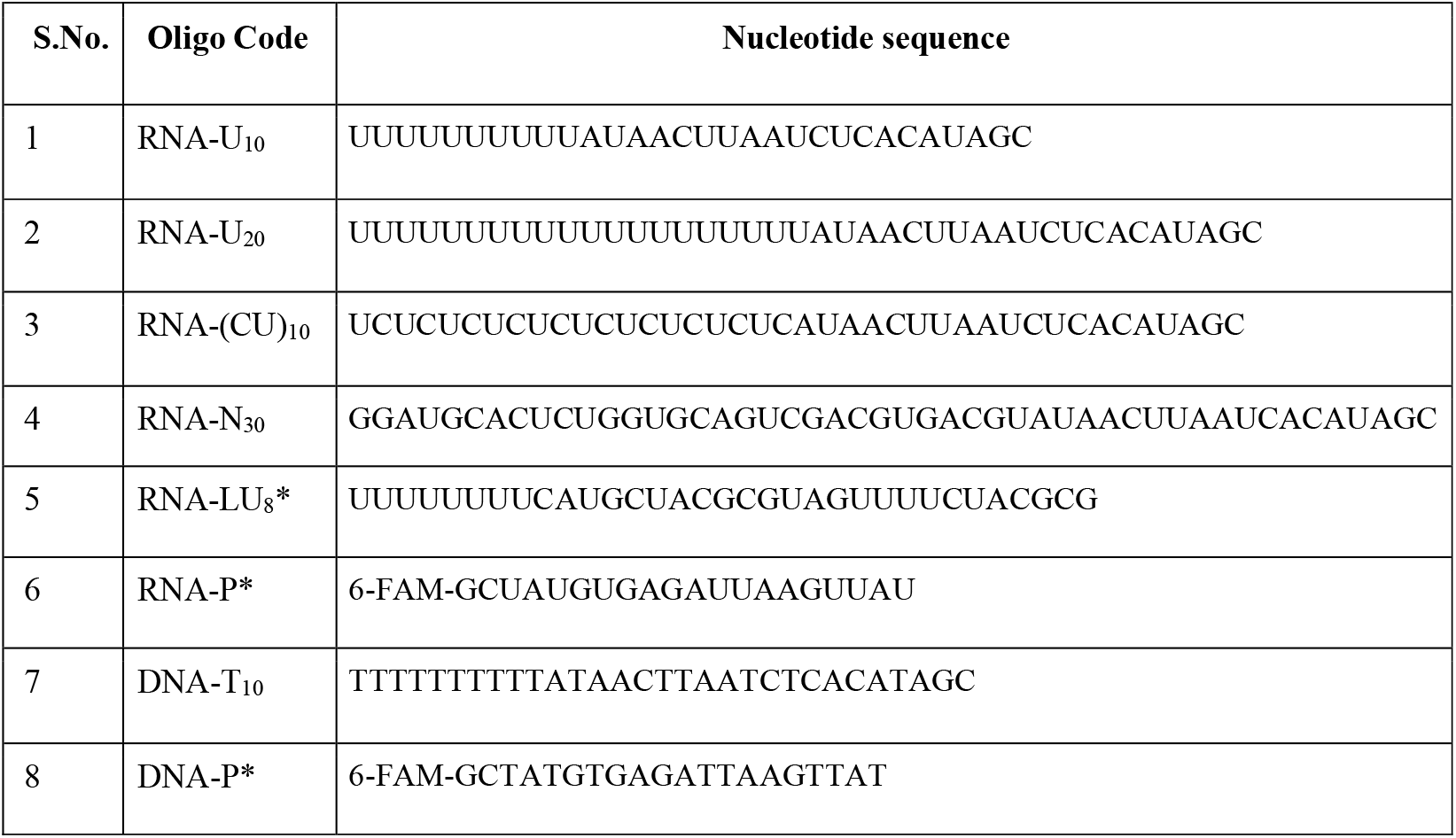
Synthetic oligonucleotides used in this study.

### 2.3. Electrophoretic Mobility Shift Assay (EMSA)

We performed EMSA to understand the interactions between RdRp holoenzyme and RNA. For this, a 20-nt 5′-6FAM-labelled RNA primer and a 30-nt RNA template having 3’ overhang of 10 uridylates were annealed as described above to produce the template-primer complex. Next, the RdRp holoenzyme was produced by mixing *NSP12, NSP8* and *NSP7* in a 1:2:1 ratio in the Polymerase Reaction Buffer (**PRB;** 20 mM Tris-HCl, pH 8.0, 10 mM NaCl, 10 mM KCl, 6 mM MgCl2, 5% glycerol, 0.01% Triton-X 100, and 1 mM DTT) and incubating overnight at 4°C (**23, 25**). To determine the optimal concentration of RdRp holoenzyme required to bind the template-primer complex, a fixed concentration of 5 μM of the template-primer complex was incubated with a series of RdRp holoenzyme concentrations (0 μM to 20 μM) in PRB for 30 minutes at 37°C and the reaction products were analysed on 10% native PAGE at 4°C. The gel was visualised using a fluorescence filter in an Azure Biosystems imaging system.

### 2.4. Fluorometric primer extension assay

Recombinant *NSP12, NSP7* and *NSP8* proteins in different combinations in the presence of different types of RNA templates or a DNA template were used to assess the polymerase activity. The primer-extension reactions were performed at 37°C using 2.5 μM *NSP12*, 2.5 μM *NSP7*, 5 μM *NSP8* and 5 μM of template-primer complex in the **PRB**. The reactions were initiated by adding 150 μM NTPs and quenched at 0 min, 30 min, 45 min, 60 min, and 120 min by adding 2X formamide gel loading buffer (**LB**; 95% formamide, 10 mM EDTA, 0.1 mg/ml xylene cyanol, and 0.1 mg/ml bromophenol blue) followed by denaturing the sample at 90°C for 10 minutes. The reaction products were resolved on 14% Urea-PAGE in 1X TBE buffer, and the fluorescence intensity of the extension products was measured at 480 nm (excitation) and 517 nm (emission) using the Azure Biosystems fluorescence filter imaging system.

### 2.5. Analysis of RNA-Protein interactions by native PAGE

To understand the molecular interactions and their consequences on the formation of the RdRp complex, interactions between *NSP12/8/7* and the RNA template in different combinations and permutations were analysed (**Table S1**). For this, two sets of experiments were performed, one set with RNA-U_10_ in complex with RNA-P* and the other with RNA-LU_8_*, the reactions were constituted in PRB in a 50 μl reaction mixture and incubated at 37°C. Briefly, the template-primer complex was pre-incubated with one of the three proteins of the RdRp complex for 15 min or 30 min and then incubated with one of the remaining two or both the components of the RdRp complex for a cumulative incubation time of 60 min. The RNA template-primer complex alone or with *NSP12* or with RdRp holoenzyme was incubated for 60 min, which served as reference controls in both the reaction setups. The reaction products, including the reference controls, were resolved on 10% Native-PAGE and the electropherograms were captured using the Fluorescence imaging system.

### 2.6. Kinetics of dissociation/association of the template-primer complex with *NSP12*

In order to understand the kinetics of dissociation of template-primer complex from *NSP12* or formation of a functional RdRp complex, we developed a method to capture the fluorescence from the 6-FAM-labelled RNA template-primer complex as well as capturing the protein profiles by Coomassie staining. For this, the preformed *NSP12*-RNA-U_10_/RNA-P* complex was incubated at 37°C with *NSP7* or *NSP 8* or both for 30-60 min and the products were resolved on 10% native-PAGE. The gels were imaged on a Fluorescence imaging system or the gel-documentation system and the fluorescent banding profiles or the protein banding profiles were captured. Next, to understand the kinetics of dissociation or formation of complexes over a period of time, the preformed *NSP12*-RNA-U_10_/RNA-P* complex was incubated with *NSP7* alone or *NSP8* alone or *NSP7/8* for up to 120 min and equal volumes of samples were collected from each reaction at regular intervals during the incubation. The products were resolved on 10% native-PAGE and the gels were imaged as described above. Thus obtained images were analysed by densitometry for quantitative estimation of dissociation/association of RNA template-primer complex.

### 2.7. Analysis of protein-protein interactions by crosslinking

Based on the 1:2:1 composition of *NSP12, NSP8* and *NSP7* in a typical RdRp holoenzyme, we sought to understand the ratios of *NSP12, NSP8* and *NSP7* when complexed in pairwise combinations. For this, 2 mg/ml stock of each *NSP* was prepared in the cross-linking buffer [**CB**; 25 mM HEPES pH 7.5, 150 mM NaCl, 5% glycerol, and 5 mM DTT] (**16**). The RdRp holoenzyme, *NSP12/8* and *NSP12/7* complexes were prepared by mixing *NSP12, NSP8* and *NSP7* in 1:2:1, 1:2 and 1:1 ratio, respectively, and incubating the reactions overnight at 4°C. Next, in order to covalently link the complexes for SDS-PAGE analysis, ethylene glycol bis (succinimidyl succinate) (EGS), prepared in DMSO, was added to a final concentration of 10 mM and the reactions were further incubated at 4°C for 2hr. Reactions were terminated by adding Tris-HCl pH 7.5 to a final concentration of 50 mM and incubating the reactions at 25 °C for 45 min. The reaction products were resolved on 8% SDS-PAGE and the electropherograms were captured.

### 2.8. Structural Modeling of *NSP12*-RNA complex with/without *NSP7* and/or *NSP8*

The structure of the RdRp complex consisting of *NSP12, NSP8*-a, *NSP8*-b, *NSP7* and RNA was retrieved from the PDB database (6YYT) (**13**). The missing regions in the *NSP12* were modelled using the SWISS-MODEL (**26**). The RNA in the cryoEM structure was replaced with the linear RNA-primer duplex (without the overhang) used in the experiments by using builder mode in PyMOL(**27**) (**Figure 1**). The modelled structures of different *NSP* complexes were subjected to MD simulations in order to understand the structural basis for the expulsion of *NSP12*-bound RNA by *NSP8* or *NSP7* when added alone, but not when added in combination. Simulations were carried out using the GROMACS software suite (version 2023) with AMBER99SB-ILDN force fields (**28**). The proteins and protein-RNA complexes were solvated in a dodecahedron box with TIP3P water model followed by neutralisation with the appropriate number of sodium or chloride ions. The neutralised system was energy-minimised using the steepest descent algorithm followed by equilibration under NVT and NPT ensemble sequentially. Production simulation was run using a leapfrog dynamic integrator with a step size of 2 fs for a timescale of 200 ns considering periodic boundary conditions in all three dimensions. Post-simulation analysis was done after eliminating periodic boundary conditions using modules available in GROMACS and *in-house* Python scripts. To check if the binding of *NSP8* or *NSP7* with *NSP12* were accompanied by changes in the RNA template/primer entry channel, the volume of the channel during the MD simulations was calculated. The volume calculations were done using ANA2 (**29**) by defining wall residues (**Table S2**) of the cavity with included_area_precision set to 1.

**Figure 1:**
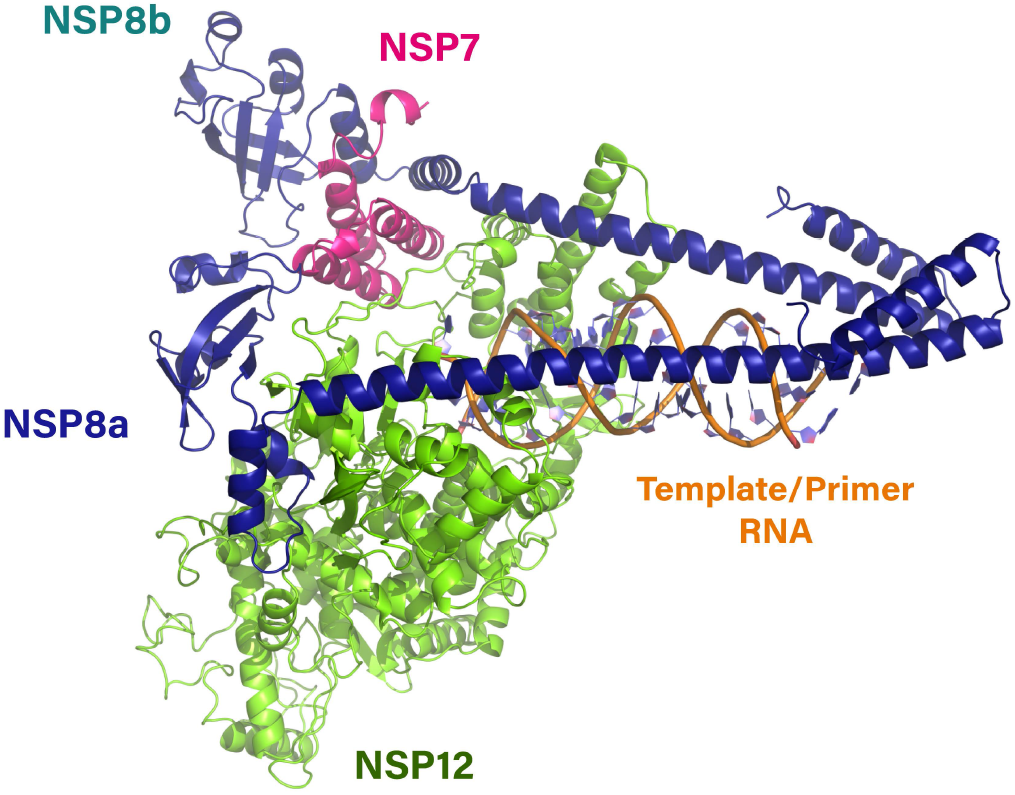
Structure of the modelled RdRp complex bound to template/primer RNA as used in experimental conditions showing *NSP12* (green), *NSP7* (magenta) and two *NSP8* monomers (navy blue and teal).

## 3. RESULTS

### 3.1. Reconstitution of SCoV-2 RdRp complex *in vitro*

The SCoV-2 RdRp holoenzyme is composed of one copy of *NSP12* interacting with one copy of *NSP7* and two copies of *NSP8*(**13, 30**). To recreate this under *in vitro* conditions, recombinant *NSP12, NSP7* and *NSP8* were produced in a bacterial expression system from their respective codon-optimised cDNA constructs, purified on Ni-NTA affinity columns followed by size-exclusion chromatography, and the same were confirmed by SDS-PAGE, and western blot assay using an anti-His antibody (**Figure 2A & 2B**). A functional RdRp holoenzyme was reconstituted *in vitro* by co-incubating *NSP12, NSP8* and *NSP7* at 1:2:1 ratio. Next, to optimise the concentration of the RdRp holoenzyme required to form the functional RdRp complex *in vitro*, 5 μM template RNA-primer (T/P*) complex (**Figure 3A**), in which the 20-nt primer was 5’-6FAM labelled, was incubated with various concentrations of the preformed RdRp holoenzyme and the products were resolved on a 10% native PAGE **(Figure 3B)**. At 0.5 μM RdRp holoenzyme, no visible RdRp complex was observed, but a visible band of the same could be seen from 1 μM onwards and the band intensities increased with increasing concentrations of RdRp holoenzyme in a dose-dependent manner (**Figure 3C**). Based on this observation, we decided to use 10 μM of the RdRp holoenzyme incubated with a 5 μM RNA template-primer (T/P*) complex (2:1 ratio) throughout this study.

**Figure 2:**
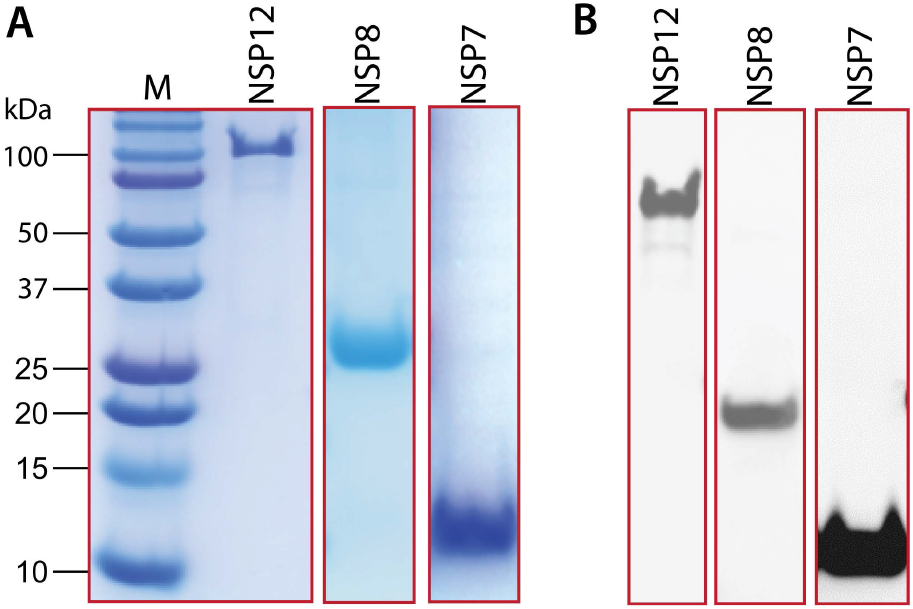
Bacterial expression of recombinant *NSP12, NSP8* and *NSP7*. **(A)** SDS-PAGE analysis of purified *NSP12, NSP8*, and *NSP7* proteins after size-exclusion chromatography **(B)** Detection of purified *NSP12, NSP8*, and *NSP7* proteins using anti-His IgG antibody.

**Figure 3:**
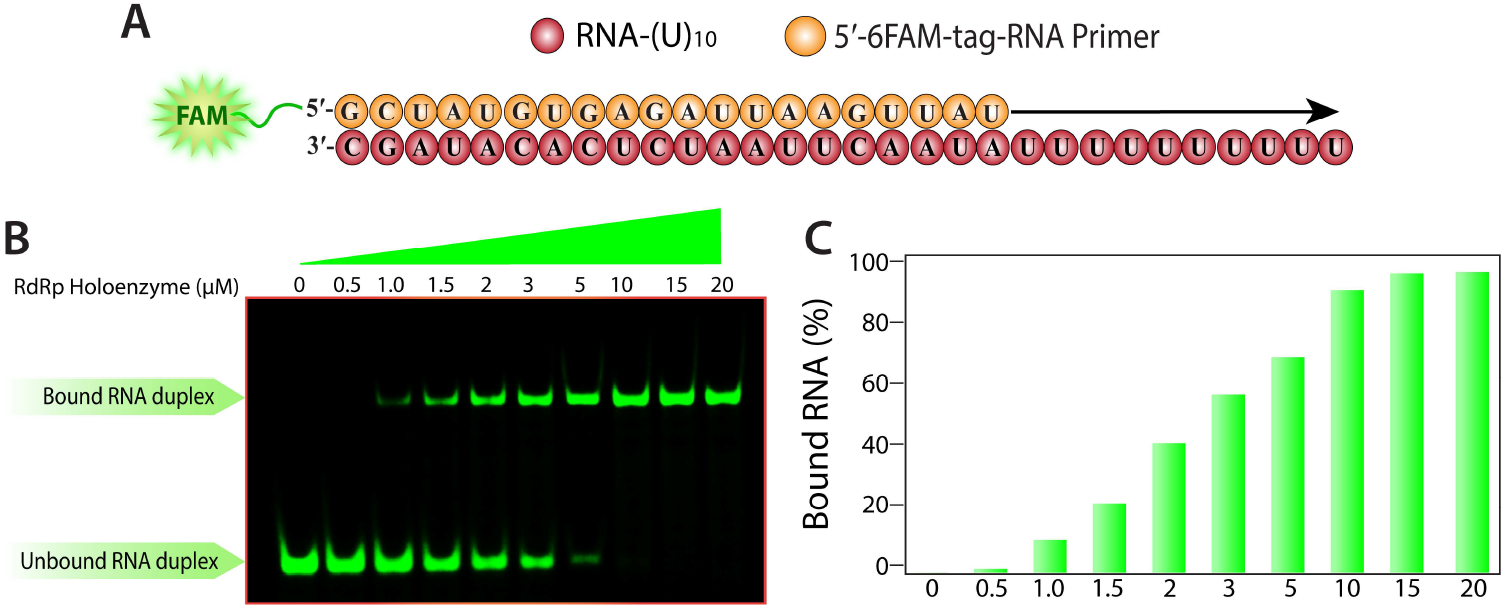
Reconstitution of SCoV-2 RdRp complex *in vitro*. **A)** Sequence of the RNA duplex with a 5′ U_10_ overhang in complex with 5′-6-FAM labelled RNA primer, **B)** Gel-based mobility-shift assay to determine the ideal concentration of the RdRp holoenzyme to achieve saturation with the RNA-Primer duplex, **C)** Quantification of bound RNA-primer complex with the RdRp holoenzyme.

### 3.2. SCoV-2 *NSP12* alone can carry out polymerase activity, which is lost when paired with *NSP7* or *NSP8*

During the coronavirus life cycle, the RdRp holoenzyme efficiently carries out the genomic RNA replication. Biochemical studies revealed that the *NSP7/NSP8* complex of SCoV-1 can perform *de novo* RNA synthesis with low fidelity on ssRNA templates, leading to the proposal that *NSP8* serves as an RNA primase to synthesise short oligonucleotide primers for the RdRp complex to catalyse complementary RNA (cRNA) synthesis (**31**). It is also well established that *NSP7* and *NSP8*, as cofactors, facilitate template RNA binding to the *NSP12* and promote the synthesis of nascent RNA (**20**). However, the individual contributions of *NSP7* or *NSP8* in RNA template binding or primer extension by *NSP12* are not clearly defined. To this end, we carried out *NSP12*-mediated primer-extension assays at 37ºC with the template-primer complex in the presence/absence of one of the two cofactors (*NSP12* alone or *NSP12*/*7* or *NSP12*/*8*) or both (*NSP12/7/8*) and the reactions were terminated at different time points post-incubation and the products were resolved on 14% urea-PAGE. To rule out template bias as well as to understand the template cooperativity with *NSP12*, we used a variety of RNA templates and a linear ssDNA template.

In these assays, the RdRp holoenzyme efficiently carried out the primer extension on linear RNA templates, irrespective of the sequence composition, in a time-dependent manner and converted almost 100% of the input primed-RNAs into full-length nascent RNA in about 45 min (**Figure 4A-D**). Its activity was marginally affected when a self-primed loop RNA was used as a template and took approximately 3-times more time to complete nascent RNA synthesis (**Figure 4E**). Interestingly, when a DNA template annealed to a DNA primer was used, it was able to incorporate dATP into the nascent strand. However, its processivity was significantly reduced and, even after 120 minutes of incubation, it could synthesise nascent DNA from only about 30% of the input DNA template **(Figure 4F**). In parallel, the activities of *NSP12* alone or in pairwise combination with *NSP7* or *NSP8*, on all the templates tested above with the RdRp holoenzyme, were analysed. Alone, *NSP12* demonstrated significant polymerase activity on all the linear RNA templates, however, it was relatively less active (30-40% less efficient) compared to the RdRp holoenzyme (**Figures 4A-D**). With regard to self-primed loop RNA, *NSP12* alone displayed significantly lower, but time-dependent, activity compared to the RdRp holoenzyme (**Figure 4E**). However, it failed to show any detectable activity with the DNA template-primer complex (**Figure 4F**). Contrary to this, when paired with *NSP7* or *NSP8*, irrespective of the template-primer complex used, *NSP12* failed to show any detectable primer-extension activity (**Figure 4A-F**). Together, these results suggest that *NSP12* alone is capable of carrying out polymerase activity, *albeit* with reduced activity compared to the RdRp holoenzyme, but its activity is completely lost when it is paired with *NSP7* or *NSP8*.

**Figure 4:**
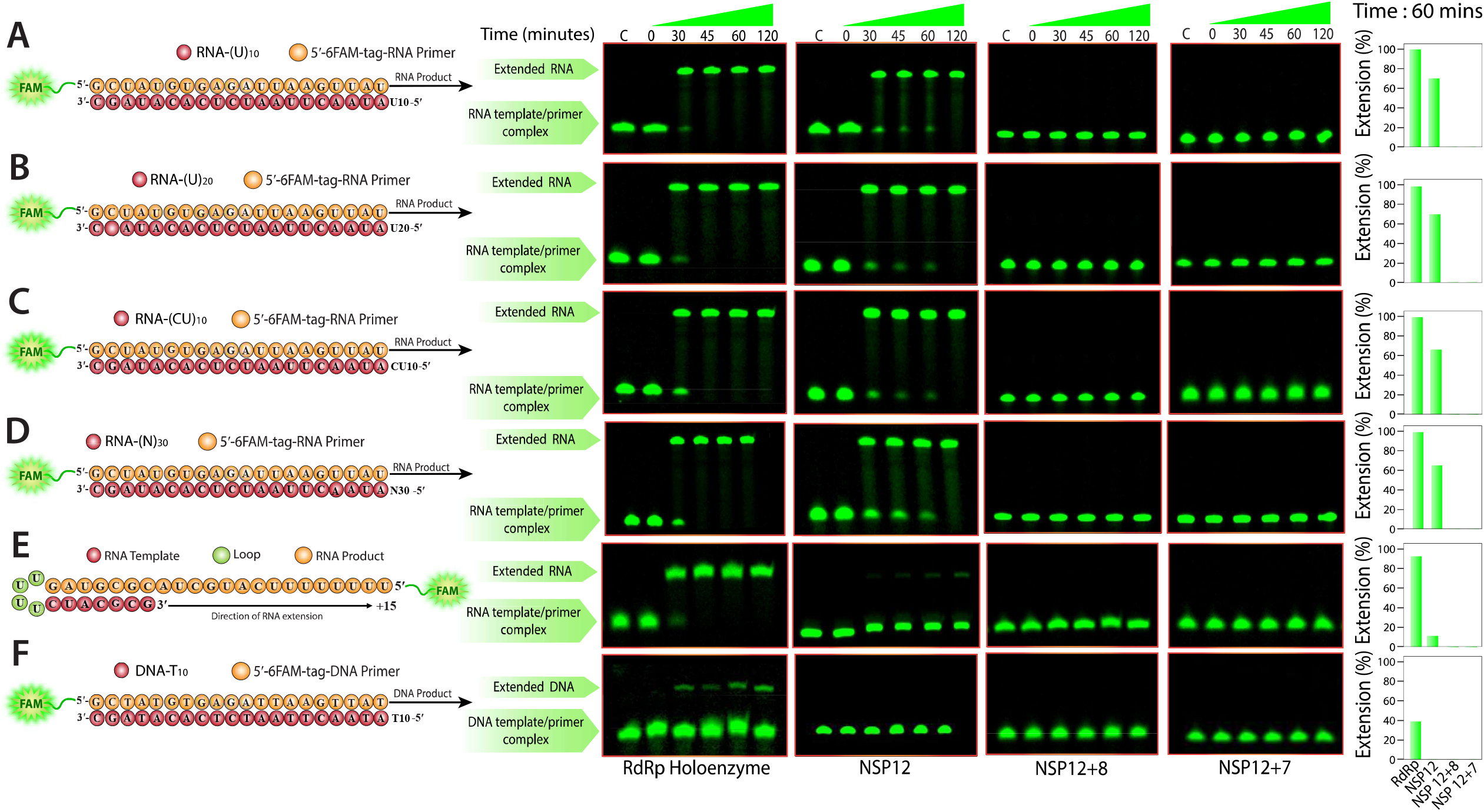
Different RNA template-primer duplexes or the DNA template-primer duplex were incubated with different *NSP* combinations and the polymerase activity was studied up to 120 min and the activities at 60 min were quantified. The quantified data is depicted in the bar graphs. **A**. RNA-U_10_, **B**. RNA-U_20_, **C**. RNA-(CU)_10_, **D**. RNA-N_30_, **E**. RNA-LU_8_*, **F**. DNA-T_10_. It may be noted that the time points and the reaction components used between different panels are the same as that used for the first panel. Accordingly, lane IDs are presented only for those in the topmost panel and the gel IDs for those in the bottom-most panel. Similarly, the bar graphs are also represented in a similar fashion.

### 3.3. *NSP7* or *NSP8* in isolation interfere with the binding of template-primer duplex with *NSP12*

In the primer-extension studies, we failed to see any detectable activity when *NSP12*/*7* or *NSP12/8* complexes were incubated with the template-primer complex. To investigate this, we analysed the interactions between the template-primer duplex and *NSP*s in different permutations and combinations, and the products were resolved on native-PAGE. Here, we also attempted to study the influence of linear template versus looped RNA on *NSP* interactions. Because it is well known that the template RNA-primer duplex readily complexes with *NSP12* or the RdRp holoenzyme, these mixtures, co-incubated for 60 min, were run alongside the test samples as known references. For this, as depicted in **Figure 5**, the template-primer complex was preincubated with *NSP12* or *NSP7* or *NSP8* or *NSP7*/*8* for 15 - 30 min followed by incubation with *NSP7* or *NSP8* or *NSP7*/*8* or *NSP12* for a cumulative incubation time of 60 min, after which the products were resolved on 10% native PAGE and visualised on a fluorescence imaging system.

**Figure 5:**
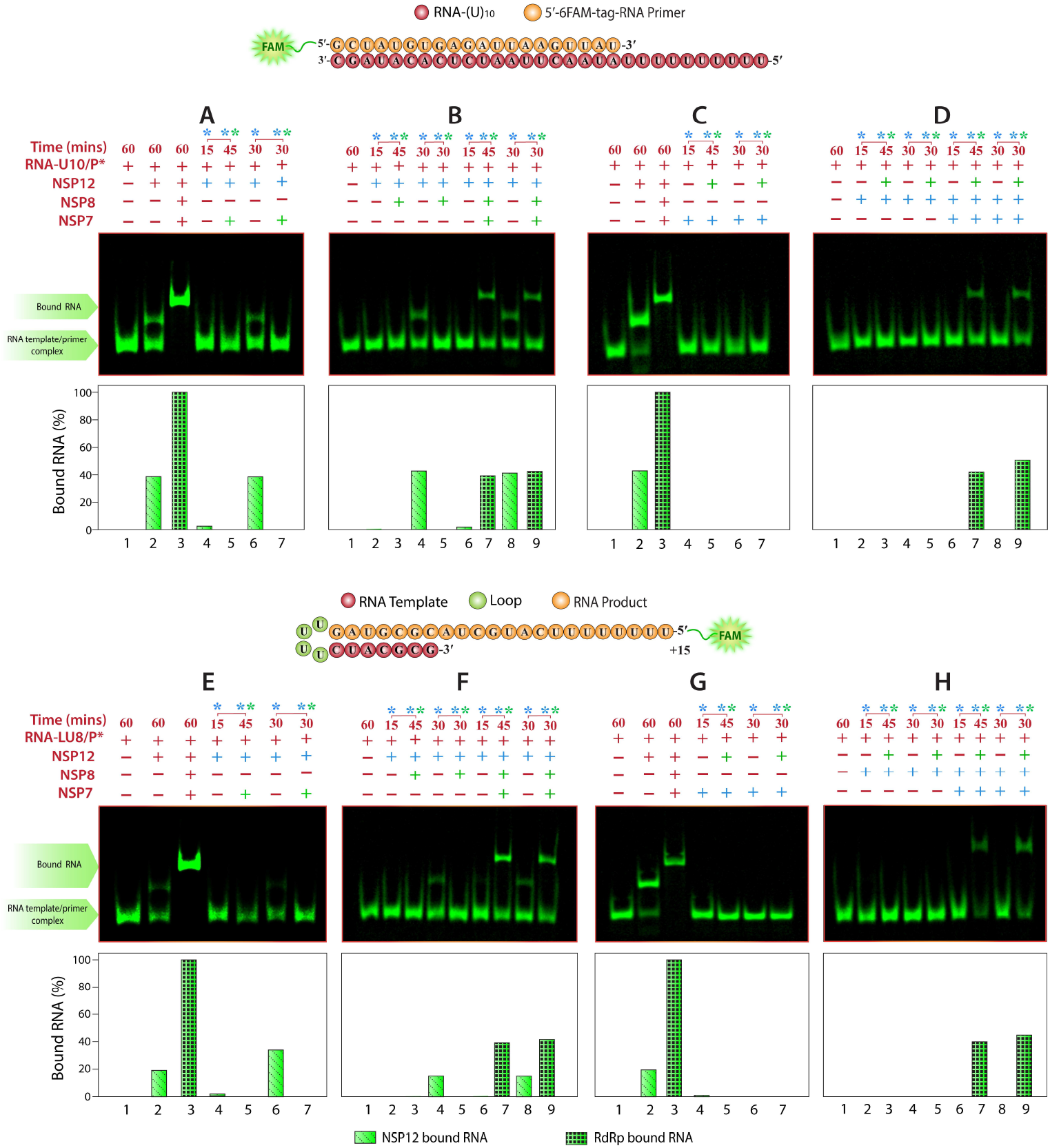
Analysis of RNA-protein interactions by EMSA. The RNA template-primer duplex was incubated with different *NSP*s for various time points, resolved on native PAGE and the fluorescent banding profiles were captured. Symbols in maroon represent the reaction components that constituted the initial reaction mixture, those in Blue represent components added first at the indicated time points during the reaction and those in green represent components added later. Bars in the plots below each gel picture represent the percentage of RNA template-primer duplex bound to different *NSP* monomers or complexes. This was calculated by subtracting the amount of fluorescence bound to *NSPs* from the total fluorescence in the lane. **A-D**. Binding reactions were carried out using the linear RNA template in complex with the fluorescently labelled primer, **E-H**. Binding reactions were carried out using the fluorescently labelled self-priming looped RNA template.

As expected, incubation of the template-primer duplex with either *NSP12* alone or the RdRp holoenzyme for 60 min resulted in stronger binding between them, which is evident from the gel mobility shift in the fluorescently-labelled RNA bands. It may be noted that the type of template had a notable influence on their interactions with *NSP12*, but not on those with the RdRp holoenzyme (**Fig. 5A, 5C, 5E & 5G, lanes 2 & 3**). When *NSP12* alone was incubated with the test RNA for 15 min, very little (quantitation data indicated a small spike) or no shift in the fluorescence, indicating an interaction, was detectable. Also, further incubation of this complex with *NSP7* or *NSP8* alone for another 45 min failed to produce the desired complexes (**Fig. 5A & 5E, lanes 4 & 5 and 5B & 5F, lanes 2 & 3**, respectively). However, when *NSP12* was incubated with the test RNA for 30 min, there was a visible *NSP12*-RNA complex, but relatively less intense compared to that incubated for 60 min. Interestingly, this complex completely disappeared upon subsequent incubation with *NSP7* or *NSP8* for an additional 30 min (**Fig. 5A & 5E, lanes 6 & 7 and 5B & 5F, lanes 4 & 5**, respectively). Contrary to this, when *NSP12* preincubated with the test RNA for 15 min or 30 min was incubated with both *NSP7* and *NSP8* for an additional 45 min or 30 min, respectively, a further shift in fluorescent bands was observed indicating the formation of the complete RdRp complex (**Fig. 5B & 5F, lanes 6-9**). Quantitatively, however, a marked increase in the band intensities of the RdRp complex, after preincubation of *NSP12* with the test RNA, was observed only when the looped template was used (**Bar graphs of Fig. 5B & 5F, lanes 8 & 9**).

Similar outcomes were observed when test RNAs were preincubated with *NSP7* or *NSP8* for different durations and then incubated with *NSP12*. Accordingly, irrespective of the time of incubation and unlike *NSP12*, when *NSP7* or *NSP8* alone was incubated with the test RNA for 15 min or 30 min, we failed to see any complex formation indicated by the absence of a shift in the fluorescent bands (**Fig. 5C & 5G, lanes 4 & 6; Fig. 5D & 5H, lanes 2 & 4**). Similarly, when these reactions are further incubated in the presence of *NSP12* for the indicated duration of time, we again failed to see any complex formation (**Fig. 5C & 5G, lanes 5 & 7; Fig. 5D & 5H, lanes 3 & 5**). Further, we also did not observe any complex formation even when both *NSP7* and *NSP8* were incubated with the test RNA for different time points post-addition (**Fig. 5D & 5H, lanes 6 & 8**). However, a clear shift in the fluorescent band was observed when these reactions were further incubated with *NSP12*, suggesting the formation of the RdRp complex (**Fig. 5D & 5H, lanes 7 & 9**). These results together suggest that the RNA templates, irrespective of their type or sequence, do not directly bind to *NSP7* or *NSP8*, but they do bind to *NSP12* when it is incubated alone with the RNA or in the presence of both the *NSP* cofactors. However, this binding between *NSP12*-RNA is absent or any existing interaction is lost when one of the two cofactors is added to the reaction. Interestingly though, the presence of both the *NSP* cofactors enhanced binding of the RNA to *NSP12* compared to that in the complete absence of either *NSP* cofactors.

### 3.4. Binding of *NSP7* or *NSP8* alone releases template-primer duplex from the *NSP12*-RNA complex

The RNA-protein binding studies suggested that *NSP7* or *NSP8* when alone, but not in combination, negatively influenced the formation of *NSP12*-RNA complexes. To gain molecular insights into the process of formation of the RdRp complex, the fate of *NSP12*-bound RNA as well as to understand the influence of the solitary presence of *NSP7* or *NSP8* on RNA binding to *NSP12*, the kinetics of these interactions were monitored by EMSA. To this end, binding reactions were performed between the RNA duplex and the three *NSP*s or between the preformed *NSP12*+RNA duplex and *NSP7* or *NSP8* or both. Replicate samples terminated at different time points post-incubation were resolved on native PAGE gels and fluorescent RNA-protein interaction profiles were captured using the fluorescence imaging system, while the Coomassie-stained gel was imaged on a gel-documentation system for protein-protein interactions. Corroborating the above findings, we found that the RNA duplex failed to bind to *NSP7* or *NSP8* alone, but bound efficiently to *NSP12* (**Fig. 6A, lanes 2-4**). Further, when the preformed *NSP12*+RNA duplex was incubated with either *NSP7* or *NSP8*, there was a decrease in the amount of *NSP12*-bound RNA duplex and a proportionate increase in the free RNA duplex (**Figure 6A, lanes 5 & 6**). As shown in **Figure 6E** (**lanes 5 & 6**), this effect was actually due to the formation of *NSP12/7* and *NSP12/8* complexes. Contrary to this, when the preformed *NSP12*+RNA duplex was incubated with both *NSP7* and *NSP8*, there was a shift in the fluorescent band whose intensity increased with time and the same is evident from the associated bar graph that represents data from the densitometric analysis of the RNA-Protein complexes (**Figure 6A, lanes 7 & 8**). Accordingly, the time-dependent formation of the RdRp holoenzyme and the intermediate complexes representing *NSP12/8* and *NSP12/7* protein-protein complexes could be seen (**Figure 6E, lanes 7 & 8**).

**Figure 6:**
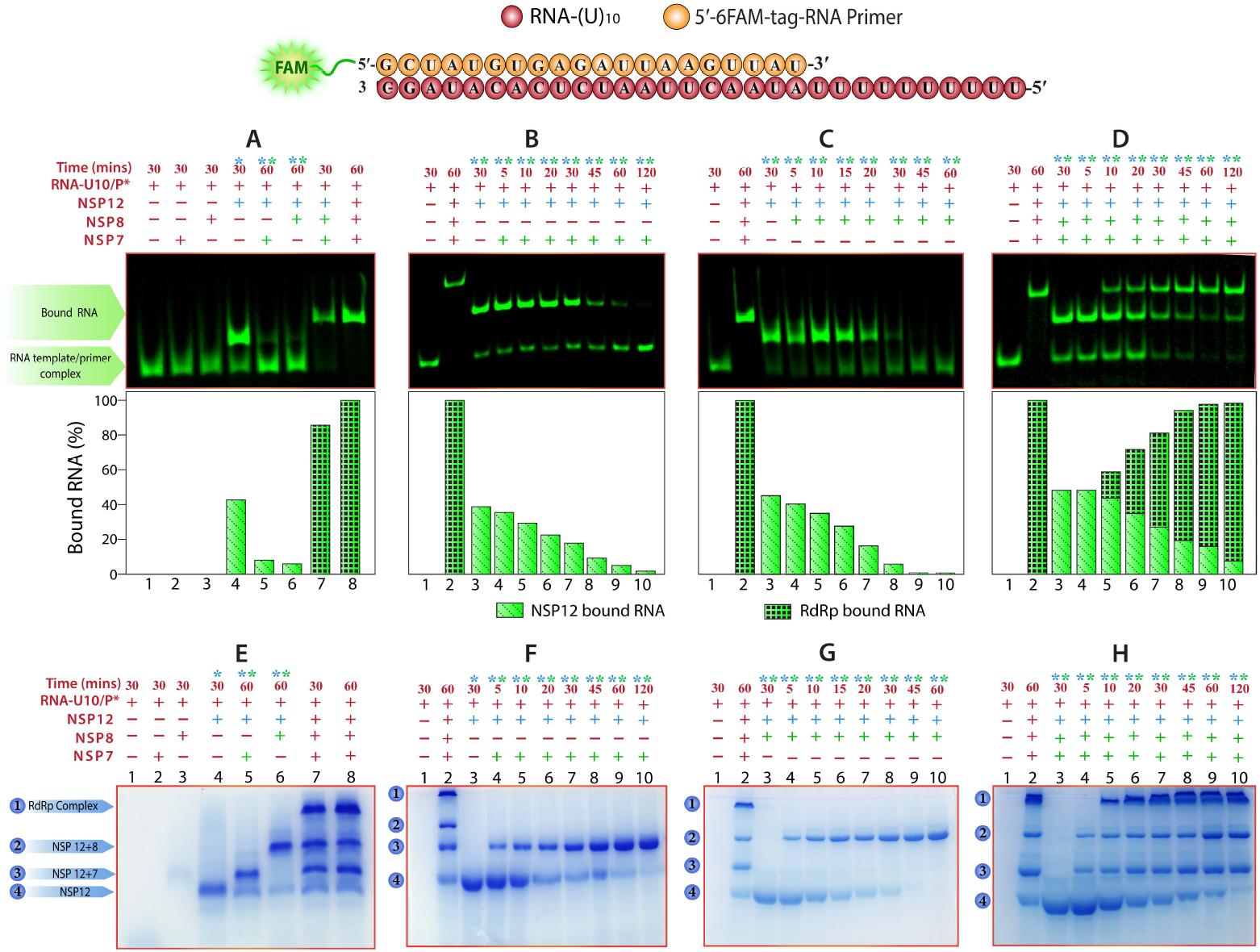
Kinetics of association/dissociation of RNA duplex from *NSP12* in the presence of either or both *NSP7* and *NSP8*. RNA duplex alone or in complex with *NSP12* were incubated with *NSP7* or *NSP8* or both, and their influence on the *NSP12*-bound RNA duplex was analysed over a period of time. Due to their small size, *NSP7* and *NSP8* ran out of the gel and could not be captured on the protein native PAGE. **A**. Binding of RNA duplex with *NSP7, NSP8* and *NSP12* were analysed and then the preformed *NSP12*-RNA complex was incubated with *NSP7* or *NSP8* or both, **B**. Kinetics of dissociation of RNA duplex from the preformed *NSP12*-RNA complex in the presence of *NSP7* alone, **C**. Kinetics of dissociation of RNA duplex from the preformed *NSP12*-RNA complex in the presence of *NSP8* alone, **D**. Kinetics of formation of the RdRp complex when the preformed *NSP12*-RNA complex was incubated with *NSP7* and *NSP8*, **E - H**. Same as ‘**A - D**’, respectively, but showing the protein-protein interactions.

In order to gain a better understanding of this phenomenon, kinetics of association/dissociation of the RNA duplex from the preformed *NSP12*/RNA duplex were investigated over a period of time in the presence of *NSP7* or *NSP8* or both. As shown in **Figure 6B and 6C** (**lanes 4 - 10**), incubation of *NSP7* or *NSP8* alone with the preformed *NSP12*-RNA complex resulted in a time-dependent decrease in the intensity of the *NSP12*-RNA complex and a proportionate increase in the intensity of the free RNA duplex. Similarly, there was a time-dependent increase in the intensity of *NSP12/7* or *NSP12/8* protein-protein complexes (**Figure 6F & 6G, lanes 4 - 10**). Notably, the dissociation of the RNA duplex was relatively faster in the presence of *NSP8* compared to that in the presence of *NSP7*. Accordingly, *NSP8* was able to displace the RNA duplex completely from the *NSP12*-RNA complex within 30 min of its addition, whereas about 5-10% of the *NSP12*-RNA complex was still intact even after 120 min of the addition of *NSP7*.

Next, the kinetics of association/dissociation of the RNA duplex from the preformed *NSP12*/RNA duplex were investigated over a period of time in the combined presence of *NSP7* and *NSP8*. Corroborating the earlier findings and contrary to that observed with *NSP7* or *NSP8* alone, an upward shift in the fluorescent band, representing the RdRp complex, was evident within 5 min of incubation, the intensity of which constantly increased over a period of time and a proportionate decrease in the intensity of the *NSP12*-bound RNA (**Figure 6D, lanes 4 - 10**). Accordingly, the densitometry-based bar graph shows a gradual decrease in the size of bars representing the unbound RNA and the *NSP12*-bound RNA, and an increase in those representing the RdRp complexes (**Fig. 6D bar graph, lanes 4 - 10)**. Similarly, the protein-protein interactions between *NSP12* and *NSP7/8* resulted in the formation of the *NSP12/7/8* complex and the concentration of the complex too increased with time (**Figure 6H, lanes 4 - 10**). Further, there was the formation of *NSP12/7* and *NSP12/8* intermediate complexes, whose concentrations increased proportionately, and a time-dependent decrease in the concentration of *NSP12* alone. Together, these results suggest that binding of *NSP7* or *NSP8* alone to *NSP12* in the *NSP12*-RNA complex displaces the RNA duplex in a time-dependent manner, and that this displacement is faster in the presence of *NSP8* compared to that in the presence of *NSP7*. On the other hand, the combined presence of *NSP7* and *NSP8* had no such effect on the bound RNA duplex, rather they facilitated the formation of the replication complex.

### 3.5. Molar ratios of *NSP7* and *NSP8* in *NSP12/7* and *NSP12/8* are similar to that in the *NSP12/7/8* complex

The EMSA data discussed above pointed towards the negative influence of *NSP7* and *NSP8*, when present alone, on *NSP12*-RNA interactions. So, prior to conducting *in-silico* analysis, to understand the consequences of the formation of *NSP12*/*7* or *NSP12*/*8* complexes on the structural organisation of *NSP12*, the assembly state and stoichiometry of these complexes were analysed by EGS-mediated protein crosslinking. Treatment of protein complexes in solution with EGS results in the formation of covalent bonds between the nearby amino acid residues through amine-reactive NHS esters (**32**). As per the published literature, the RdRp holoenzyme is estimated to be an approx. 156 kDa complex, in which *NSP12, NSP8* and *NSP7* are assembled in a 1:2:1 ratio (**13**). As shown in **Figure 7**, EGS-mediated crosslinking of the three *NSP* mixtures produced the RdRp holoenzyme of the expected size. Next, incubation of *NSP12* with *NSP8* alone or *NSP7* alone under similar reaction conditions followed by EGS-mediated crosslinking produced complexes with approx. weights of 146 kDa and 110kDa, respectively, suggesting that the *NSP12/8* complex is composed of one *NSP12* monomer (100kDa) and two molecules of *NSP8* (46kDa), while the *NSP12/7* complex comprises one *NSP12* monomer (100kDa) and one *NSP7* monomer (10kDa) **(Figure 7)**. Crosslinking of *NSP8* alone, *NSP7* alone, and *NSP8*/*7* mixtures yielded complexes similar to those reported in the literature (data not shown) (**16**). Together, the above data provided critical information pertaining to the molar ratios of the *NSP* monomers in different complexes in solution, and the same was used to constitute the complexes for computational studies.

**Figure 7:**
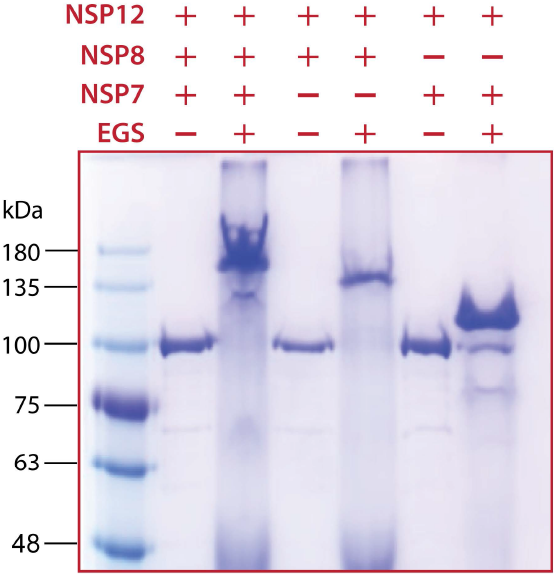
SDS-PAGE showing RdRp, NSP12/8, and NSP12/7 complexes in the presence and absence of crosslinker (EGS).

### 3.6. Binding of *NSP7* or *NSP8* alone to *NSP12* leads to constriction of the RNA entry/binding tunnel

Previous studies on the roles of SCoV-2 *NSP*s in viral genome replication and transcription have attributed the polymerase activity to *NSP12* and proposed that the association of *NSP12* with *NSP7* and *NSP8* cofactors is mandatory for this activity (**33**). However, the molecular interplay between them and the influence of each cofactor on RNA binding and replication remains largely unknown. In the present study, the *in vitro* biochemical data suggested that both RdRp holoenzyme and *NSP12* exhibit specific RNA-binding and polymerase activities, *albeit* with varied efficacies. Further, the data also suggested that binding of *NSP8* or *NSP7* alone to *NSP12* prevents RNA binding or results in the dissociation of bound RNA from *NSP12*. To understand the mechanism underlying the *NSP8* and *NSP7* mediated regulation of the *NSP12*-RNA interaction, MD simulations were performed for 200 ns for each of the *NSP* combinations tested in the *in vitro* biochemical studies and the volume of RNA binding/entry tunnel in *NSP12* in each complex was calculated using ANA2 by defining the wall residues of the tunnel (**Figure 8A)**. The tunnel volumes of all complexes are given in **Table S3**. The *NSP* ratios observed in the cross-linking experiments were used to constitute the complexes for the analysis. Since the RdRp complex is missing the electron density at several places, the model of the RdRp holoenzyme in complex with RNA was modelled and deposited on the ModelArchive database (ID-ma-c0hnm).

**Figure 8:**
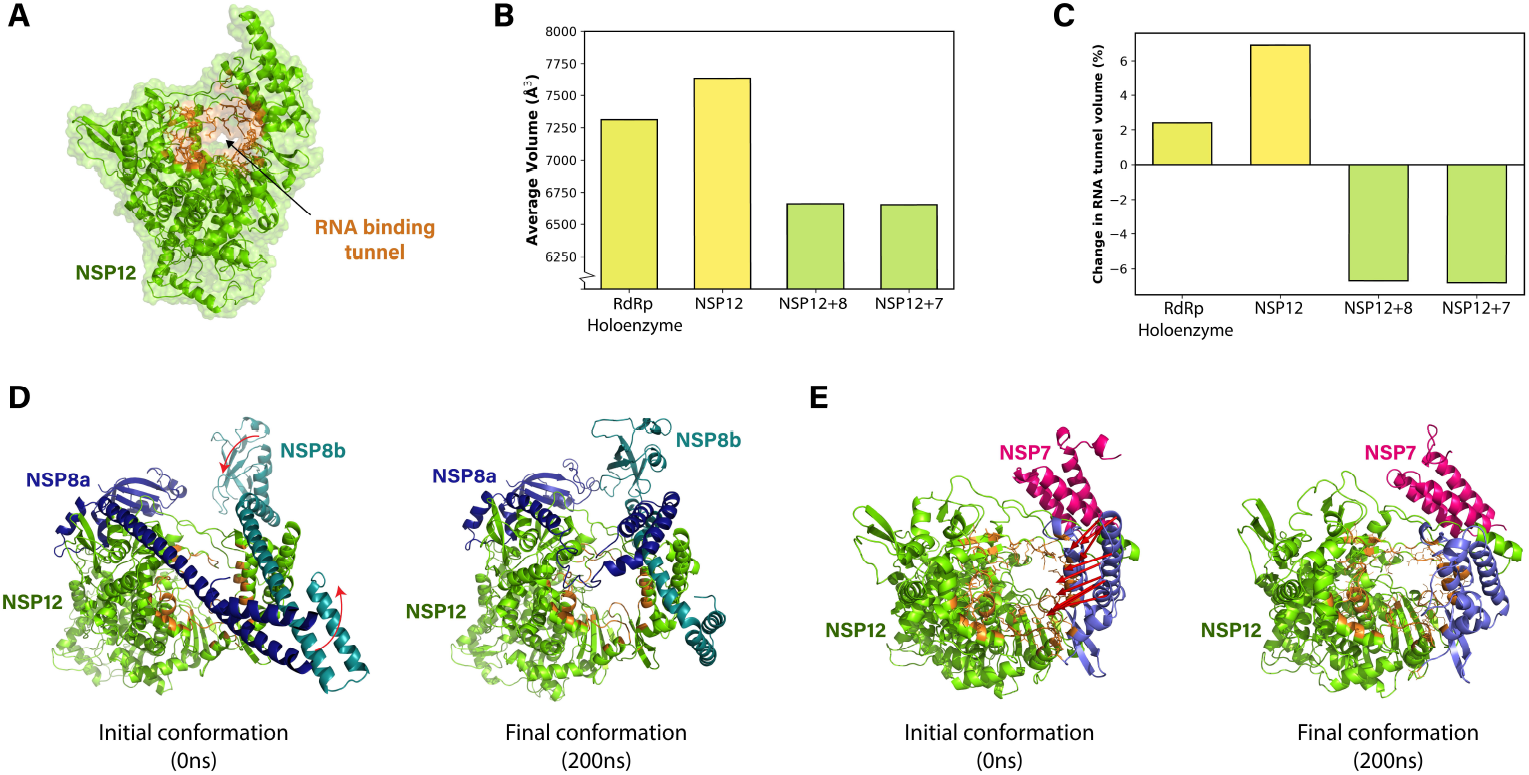
MD simulations analyses of different *NSP* complexes and calculation of the RNA binding/entry tunnel volume. **(A)** Structure of *NSP12* (green) highlighting the residues forming the RNA binding tunnel (orange) selected as wall residues for the volume calculations. **(B)** Bar graph showing an average volume of the RNA binding tunnel as calculated for the 200ns simulation using ANA2 **(C)** Bar graph showing percentage changes in the average volume of the RNA-binding tunnel compared to the average volume of RNA-binding tunnel in the RdRp complex calculated after removal of the RNA in post-processing of the trajectory. **D**. Initial and final conformations of *NSP12/8* complex during the simulation shown in cartoon representation. The arrows indicate *NSP8-a* bending and interaction with *NSP8-b* and the thumb domain, **E**. Structure of the initial and final conformations of *NSP12/7* complex with finger domain shown in blue. During the simulation, the finger domain of *NSP12* moved towards the RNA binding (shown in orange) tunnel as depicted by red arrows, which show displacement greater than 10 Å.

The RdRp complex comprising the RNA bound to the RdRp holoenzyme was studied as the reference control and the structure of *NSP12* was stable without any no notable changes during the MD simulations analysis (**Figure 8B**). Similar to the RdRp complex, RNA bound to *NSP12* too was stable without showing any notable changes, but the RNA duplex was a little bent most likely due to a lack of support in the absence of *NSP8* monomers. When the RdRp holoenzyme alone, without the RNA, was analysed, both the *NSP8* monomers, *NSP8*-b and *NSP8*-a, displayed significant movement in the N-terminal helical extensions. Accordingly, the N-terminal helical extension of *NSP8*-b folded backwards and interacted with itself and the tip of the *NSP12* thumb domain, whereas the pi-helix (Lys58-Arg75)-harbouring *NSP8*-a lost its secondary structure through the extension of its N-terminus and became predominantly solvent exposed. The mean volume of the RNA binding tunnel in the RdRp complex, after removing the RNA, was found to be 7560 Å^3^, whereas it was only marginally higher (7734 Å^3^) for *NSP12* alone.

Contrary to this, significant rearrangements were observed in the conformation of *NSP12* when the *NSP12/8* complex was analysed. Accordingly, in the absence of *NSP7*, the C-terminus of *NSP8*-b slid into the void space, which is otherwise occupied by *NSP7*, and interacted with the finger extension of *NSP12* and the C-terminus of *NSP8*-a. Further, the N-terminal helical extension of *NSP8*-b wrapped the base of the thumb domain, while the *NSP8*-a was bent allowing its N-helical extension to interact with itself and the tip of the *NSP12* thumb domain, thus making the complex more compact (**Figure 8D**). These conformational changes hugely impacted the volume of the RNA entry tunnel and the same was found to be 6790 Å^3^ (**Figure 8B**). When the *NSP12*/*7* complex was analysed, though no major conformational changes were observed in the *NSP12*, the thumb domain of *NSP12*, similar to that observed for the *NSP12/8* complex, showed a movement towards the RNA binding region leading to constriction of the RNA entry/binding tunnel (**Figure 8E**). Further, the finger extension, which connects the thumb and the finger domain, was also found to be retracted and therefore contributed to the observed constriction of the RNA entry/binding tunnel. Accordingly, the mean volume of the RNA entry tunnel in the *NSP12/7* complex was calculated to be 6831 Å^3^ (**Figure 8B**). Together, the above data suggested that binding of *NSP7* or *NSP8* alone to *NSP12* resulted in a decrease in the RNA binding/entry tunnel volume of *NSP12* by about 650 - 700 Å^3^ compared to that in the RdRp complex. Overall, the comparison of *NSP12/8* and *NSP12/7* with the RdRp complex showed a decrease of 6.7 and 6.8% in the RNA binding tunnel volume, respectively, whereas RdRp holoenzyme and *NSP*12 alone showed an increase of 2.4 and 6.9%, respectively (**Figure 8C**). Since *NSP12/8* and *NSP12/7* complexes did not allow RNA template/primer binding with the NSP12, we could not perform the MD simulations analysis for *NSP12/8*/RNA and *NSP12/7*/RNA complexes. The structural changes observed in *NSP12, NSP8-a, NSP8-b* and *NSP7*, in different complexes during the simulations, were calculated in terms of RMSD, RMSF and radius of gyration (**Figure S2, S3, S4** and **S5** respectively).

## DISCUSSION

### Individual contributions of the *NSP* cofactors are poorly understood

RNA-dependent RNA polymerase is a critical component of the RNA virus machinery, and in coronaviruses, *NSP12*, along with its cofactors *NSP7* and *NSP8*, carries out this activity. Currently approved CoV RdRp inhibitors, such as nucleoside analogs, have been found to offer limited clinical benefits, and the presence of an exonuclease in the viral machinery further leads to the development of resistance to these nucleoside analogs (**34**). This scenario necessitates the identification of alternate therapeutic options in the viral replication machinery so as to develop more effective therapies. In this direction, there is contrasting data, mainly from studies on SCoV-1 and SCoV-2, on the ability of *NSP12* alone to carry out RNA catalysis in the absence of the two cofactors. Some have projected that the two cofactors are indispensable for *NSP12*-mediated viral RNA replication, while others have shown notable RNA polymerase activity for *NSP12* in the complete absence of the two cofactors (**19, 20**). Available literature suggests critical roles for *NSP7* and *NSP8* cofactors in facilitating template RNA binding to and in enhancing the activity of *NSP12*. However, the mechanism by which they facilitate these activities and their individual contributions to these activities remain unexplored. Thus, in order to exploit the therapeutic options they may offer, it is imperative to understand the architecture and dynamics of *NSP7/8* interactions with *NSP12*.

### The activity of the RdRp complex is independent of the template sequence but is influenced by the structure

In the present study, using the bacterially expressed recombinant *NSP*s, we could successfully reconstitute a functional RdRp complex *in vitro* by mixing *NSP12, NSP8* and *NSP7* in 1:2:1 ratio to produce the RdRp holoenzyme and incubating the complex with a linear RNA template bound to a fluorescently-labelled primer. The *in vitro* reconstitution of the RdRp complex occurred in a dose-dependent manner, and a template-RdRp holoenzyme ratio of 1:2 was found to be optimal for RNA binding (**Figure 3**). Next, in order to investigate the processivity of *NSP12* and the influence of each of the two *NSP* cofactors on processivity, primer extension assays were performed using a variety of templates, including a self-priming looped RNA and a linear DNA template bound to a DNA primer (**Figure 4**). Our data suggest differential processivity of the RdRp holoenzyme on linear and looped templates. Although the holoenzyme was equally active, irrespective of their sequence composition, on a variety of linear RNA templates, its activity on a looped RNA template was at least 3-times lower compared to that on a linear template. These results with the linear and looped RNA templates suggest that the looped RNA templates probably pose certain structural limitations and therefore may not be ideal for RdRp activity studies.

Further, the holoenzyme exhibited detectable, but significantly slower, DNA-dependent-DNA-polymerase activity on a linear DNA template/primer complex (**Figure 4F**). This is the first instance of a DNA-dependent-DNA polymerase activity reported for a coronavirus RdRp holoenzyme, which is not entirely surprising, as a previous study has reported such an activity, but using SCoV-1 *NSP12* alone (**19**). Similarly, Cheng et.al. also reported weak RNA-dependent-DNA polymerase activity in their filter-binding polymerase assays using SCoV-1 *NSP12* alone (**35**). These studies highlight some yet unknown differences between the RdRp holoenzymes of SCoV-1 and SCoV-2.

### SCoV-2 *NSP12* alone is capable of carrying out efficient RNA catalysis

With regard to *NSP12*, there is contrasting data from SCoV-1 and SCoV-2 studies as mentioned earlier. Importantly, the majority of studies on SCoV-2 *NSP12* used self-priming looped RNA as the template in their primer-extension assays. Contrary to the existing data on SCoV-2 *NSP12*, our data clearly suggested that *NSP12* alone can carry out the efficient primer-dependent extension of RNA, although the rate of its activity was approximately 25% slower on linear RNA templates and about 90% slower on a looped RNA template compared to the RdRp holoenzyme (**Figure 4**). Notably, the prevalent use of loop templates in SCoV-2 studies misled previous studies to conclude that *NSP12* alone is not active in the absence of the cofactors (**13, 25, 36**). With regard to its ability to catalyse nascent DNA synthesis from the DNA primer on a DNA template, we failed to see any detectable primer extension activity, even after longer incubations beyond 120 min. This relatively slower catalysis of RNA synthesis from linear RNA templates, highly inefficient catalysis from looped RNA templates, and lack of DNA catalysis for *NSP12* compared to the RdRp holoenzyme points towards the presence of unfavourable conformations in the template binding tunnel in *NSP12*. Further, this also highlights the potential role of *NSP7* and *NSP8* in creating favourable conditions for efficient template binding to and/or facilitating proper interactions between catalytic site residues in *NSP12*. Together, a comparison of the polymerase activities of the RdRp holoenzyme and *NSP12* suggest compromised processivity in the absence of *NSP7* and *NSP8*.

### *NSP7* and *NSP8*, when alone, affect *NSP12* interactions with template RNA

While the RNA binding and polymerase activities of *NSP12* in the presence of both *NSP* cofactors are well described in the literature, the polymerase activities of *NSP12* in complex with *NSP7* or *NSP8* have not been well explored. To this end, we failed to see any detectable polymerase activity for the *NSP12/7* or *NSP12/8* combinations in our primer extension studies (**Figure 4**). So, we next carried out EMSA studies, wherein we found that the *NSP12/7* and *NSP12/8* complexes failed to bind to the RNA duplex (**Figure 5**). Similar results were obtained when *NSP12* was incubated with the RNA duplex before or after the addition of the individual *NSP* cofactors. Further, the addition of *NSP7* or *NSP8* to the preformed *NSP12*-RNA complex and the kinetics of their effects on the RNA duplex were monitored, and we found that both combinations caused the release of the bound RNA duplex in a time-dependent manner, but this dissociation was notably faster in the presence of *NSP8* (**Figure 6B & 6C**). The protein-protein interaction data further supported that the observed effects on the RNA duplex were actually due to the formation of the *NSP12/7* and NSP12/*8* complexes (**Figure 6F & 6G**). However, when both the cofactors were added to the preformed *NSP12*-RNA complex, we found that the RdRp complexes were formed in a time-dependent manner (**Figure 6D & 6H**). Taken together, these results suggest that the presence of only one of the two cofactors alters the conformation of *NSP12*, which is not ideal for RNA duplex binding.

### Regulatory role of *NSP8* and *NSP7* proteins in *NSP12*-RNA interaction

Next, we set out to understand the regulatory roles of the two cofactors, precisely pertaining to the *NSP12-*RNA interactions, through *in silico* studies. For this, it was important to analyse the number of copies of the respective cofactors in the *NSP12/7* and *NSP12/8* complexes. In this regard, our protein crosslinking experiments confirmed that the stoichiometry of the *NSP12/7* and *NSP12/8* complexes is the same as that of the RdRp holoenzyme **(Figure 7)**. We then used this information to generate atomistic models of the two complexes along with other reference complexes. Using these models, *NSP12*-specific structural changes that influence the binding of the RNA duplex to *NSP12* were investigated using MD simulation analyses **(Figure 8)**. Compared with the RdRp complex, no significant changes were observed in the *NSP12* of the RdRp holoenzyme, and the mean volume of the RNA entry tunnel increased marginally by 2.4%. With regard to *NSP12* alone, though no notable structural changes were observed in the MD simulations, there was a notable increase of 6.9% in the tunnel volume. It may be assumed that such a significant deviation in the tunnel volume compared to that in the RdRp complex may not be ideal for optimal binding of the template RNA, which would also expectedly affect the processivity of the enzyme as well. This probably explains the relatively slower RNA catalysis on linear RNA templates by *NSP12* alone or its poor activity on looped RNA substrates and the lack of DNA-dependent DNA polymerase activity.

On the other hand, due to its high conformational flexibility, *NSP8* in the *NSP12/8* complex adopts a compact conformation through intermolecular interactions between *NSP8-a, NSP8-b* and *NSP12*. As a result, the thumb domain of *NSP12* moved towards the tunnel, thereby constricting the RNA-binding cavity by 6.7% compared to that in the RdRp complex and making the same highly unfavourable for RNA binding **(Figure 8D)**. Additionally, because of the extensive interfacial interactions between *NSP8* and *NSP12, NSP8* has the potential to induce rapid conformational changes in *NSP12*. Although the conformation of *NSP12* did not change much in the *NSP12/7* complex, the thumb domain of *NSP12*, similar to that seen in the *NSP12/8* complex, moved towards the tunnel, thereby significantly reducing the tunnel volume by 6.8% **(Figure 8E)**. Together, compared to the RdRp complex, the volume of the RNA binding tunnel decreased by approximately 650 Å^3^ in the *NSP12/8* and *NSP12/7* complexes, which is very significant and thus contributes to the lack of RNA binding to *NSP12* and/or release of the bound RNA duplex from the preformed *NSP12*/RNA complexes. Notably, while both *NSP8* in the *NSP12/8* complex and *NSP7* in the *NSP12/7* complex inhibited the *NSP12*-RNA interaction through the constriction of the RNA-binding tunnel, *NSP8* may also impose steric obstruction. Detailed structural studies of these complexes may shed more light on the mechanistic details of this inhibition.

### Conclusions drawn from the study

Finally, to conclude, using the bacterially expressed recombinant SCoV-2 *NSP12, NSP7* and *NSP8*, we have reconstituted a functional RdRp complex capable of efficient primer-dependent RNA catalysis independent of type and sequence of the template. Additionally, it also displayed DNA-dependent-DNA polymerase activity, which was significantly slower compared to its RdRp activity. While *NSP12* alone too demonstrated the primer-dependent RdRp activity, its activity was relatively slower on linear RNA templates compared to the RdRp complex, whereas it demonstrated very poor RNA catalysis on the self-priming looped RNA template and no DNA-dependent DNA synthesis. Together, these results reemphasize the existing hypothesis that *NSP7* and *NSP8* play an important role in RNA replication. As part of our attempts to delineate the individual contributions of *NSP7* and *NSP8*, precisely in facilitating template RNA binding to *NSP12*, we have clearly demonstrated that *NSP12*, in pairwise combination (*NSP12/7 & NSP12/8*), fails to bind to the template RNA and therefore lacks polymerase activity. Indeed, the presence of *NSP7* or *NSP8* in complex with *NSP12* induced the release of any previously bound RNA in a time-dependent manner. Later, through *in silico* MD simulation analysis, we demonstrated that the lack of RdRp activity for *NSP12/7* and *NSP12/8* is due to the *NSP7*- and *NSP8*-induced structural changes in *NSP12*, which led to severe constriction of the RNA entry/binding tunnel, thereby creating unfavourable conditions for RNA entry/binding. Contrary to this, relatively weak polymerase activity of *NSP12* on different templates was due to the presence of an increased RNA entry/binding tunnel volume compared to that in the RdRp holoenzyme, which does not allow optimal binding of the template to allow efficient polymerase activity. Targeted structural studies are required to further understand the mechanistic details of the observed inhibition for different *NSP* combinations.

## AUTHOR CONTRIBUTIONS

DS, TK, RK, MBA & KKI: conceived and designed the experiments; DS, TK, RK, VK, KB, SHT, AG, MRK, MC, SRK: Methodology; TK, MK, AP & KKI; Software; DS, TK, RK, DS, MBA & KKI: Data analysis; DS, TK, RK, SK, MRK, SG & MC: Investigation and data curation; DS, TK, RK, SHT, SG, MC, MBA & KKI: Writing – original, draft, review and editing; MBA & KKI: Project administration; KKI: Funding acquisition. All authors have read and agreed to the published version of the manuscript.

