## Supporting Information for "Interaction of SCoV-2 *NSP7* or *NSP8* alone with *NSP12* causes constriction of the RNA entry channel: Implications for novel RdRp inhibitor drug discovery"

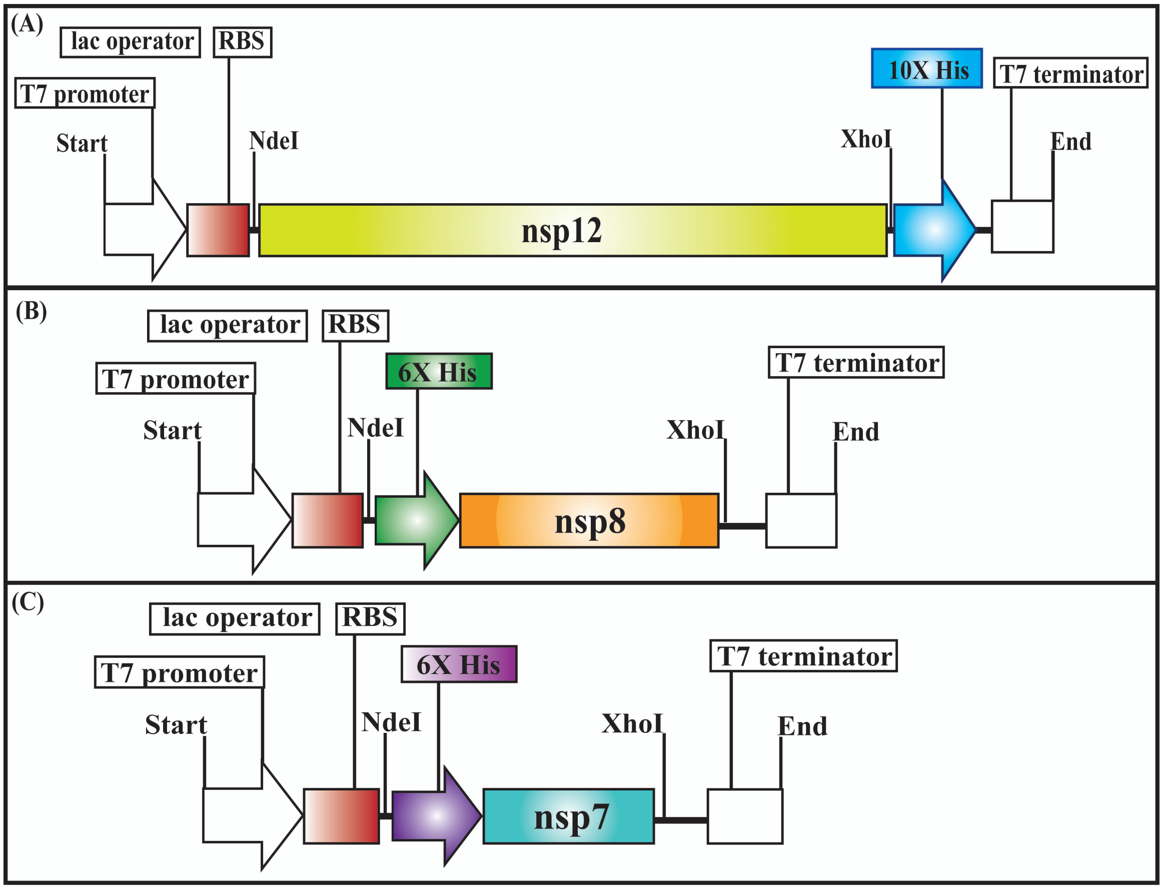


**Figure S1** illustrates the cloning strategies in the study. The codon-optimized and custom-synthesized cDNA sequences were cloned into pET22b (*NSP12*), pET28a (*NSP8* & *NSP7*) vector system, and the constructs were used to transform *E. coli* BL21(DE3) cells for expression studies. **A - C.** Strategies used to clone the cDNAs coding for SCoV-2 *NSP12*, *NSP8* and *NSP7*.

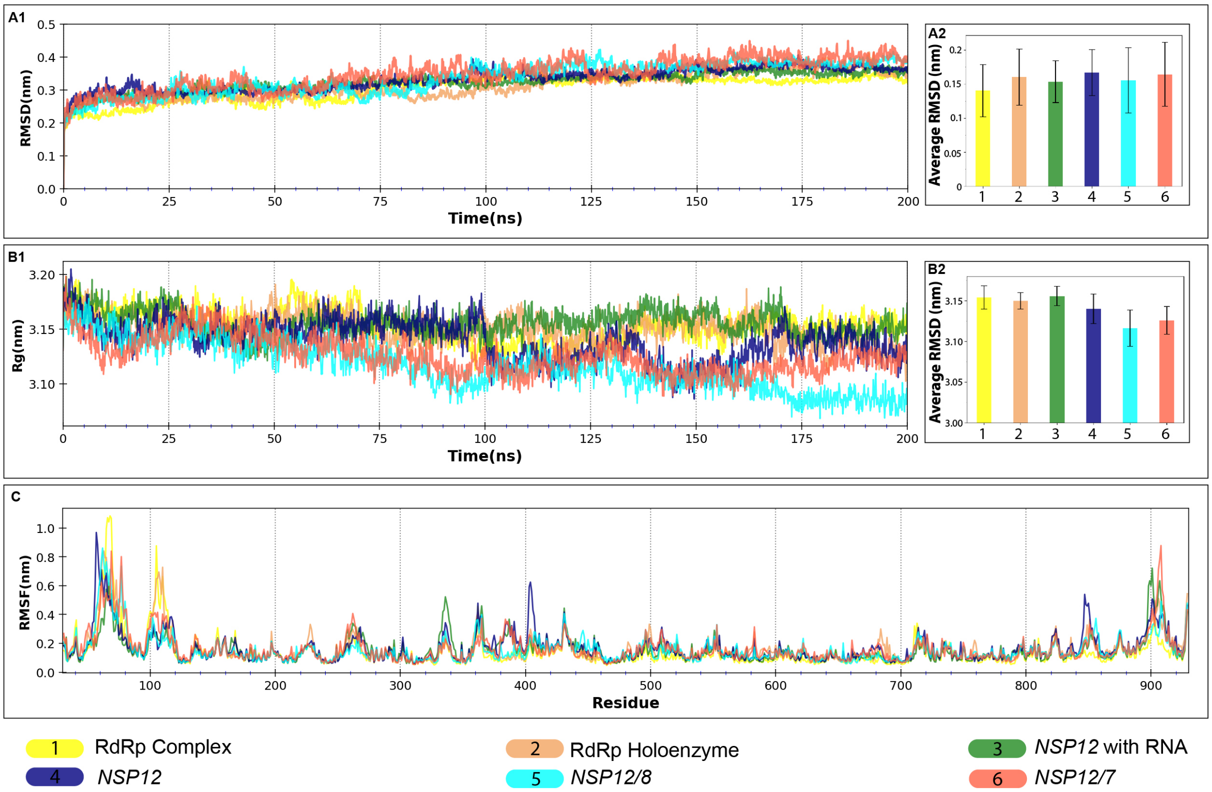


**Figure S2** showing the MD simulations analyses for *NSP12* in different complexes. **A1.** RMSD, **B1.** Rg, **C.** RMSF. The bar charts **A2** and **B2** placed adjacent to **A1** and **B1** illustrate the statistical representation of the data.

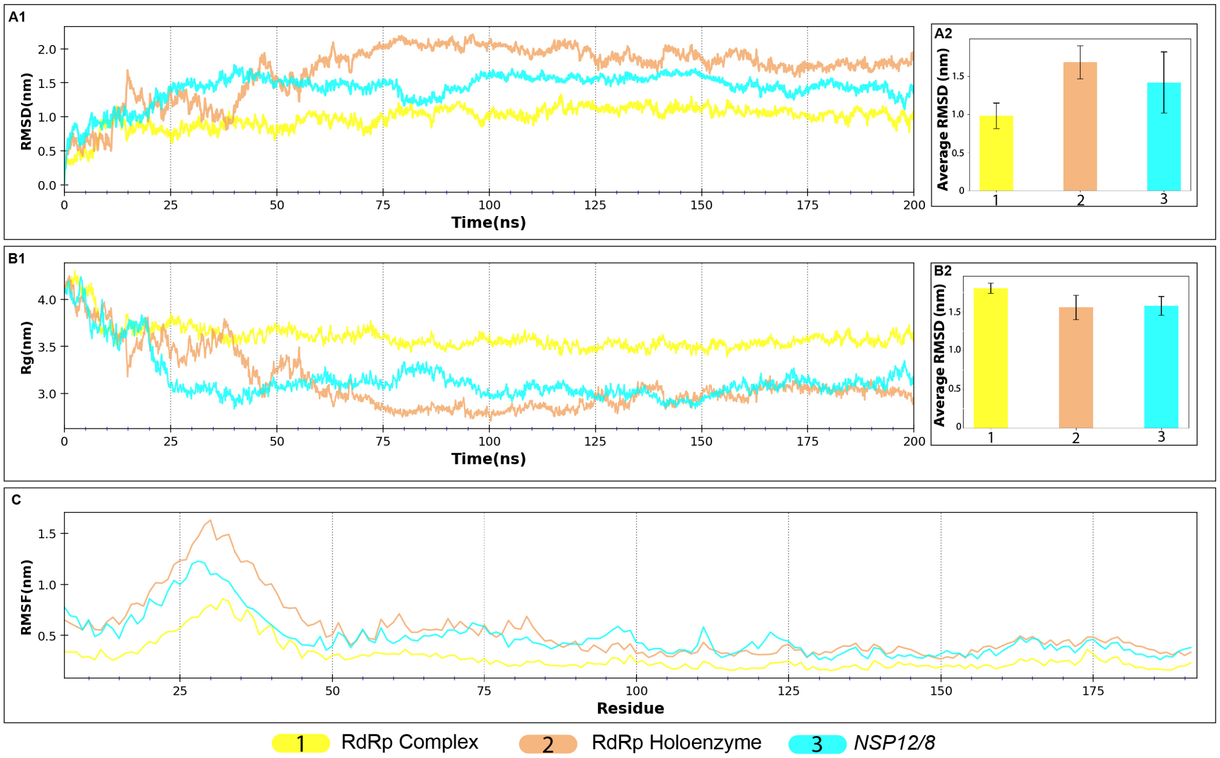


**Figure S3** showing the MD simulations analyses for *NSP8a* in *NSP12/8* complex. **A1.** RMSD, **B1.** Rg, **C.** RMSF. The bar charts **A2** and **B2** placed adjacent to **A1** and **B1** illustrate the statistical representation of the data.

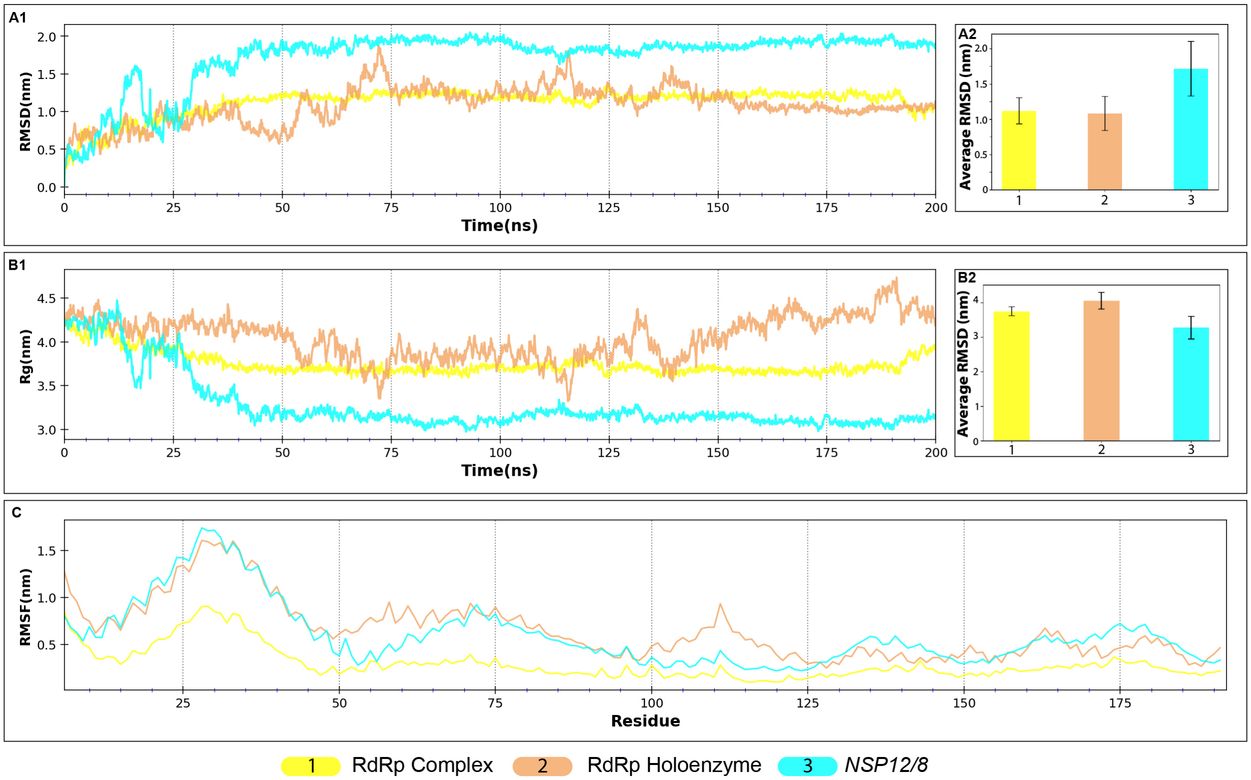


**Figure S4** showing the MD simulations analyses for *NSP8b* in *NSP12/8* complex. **A1.** RMSD, **B1.** Rg, **C.** RMSF. The bar charts **A2** and **B2** placed adjacent to **A1** and **B1** illustrate the statistical representation of the data.

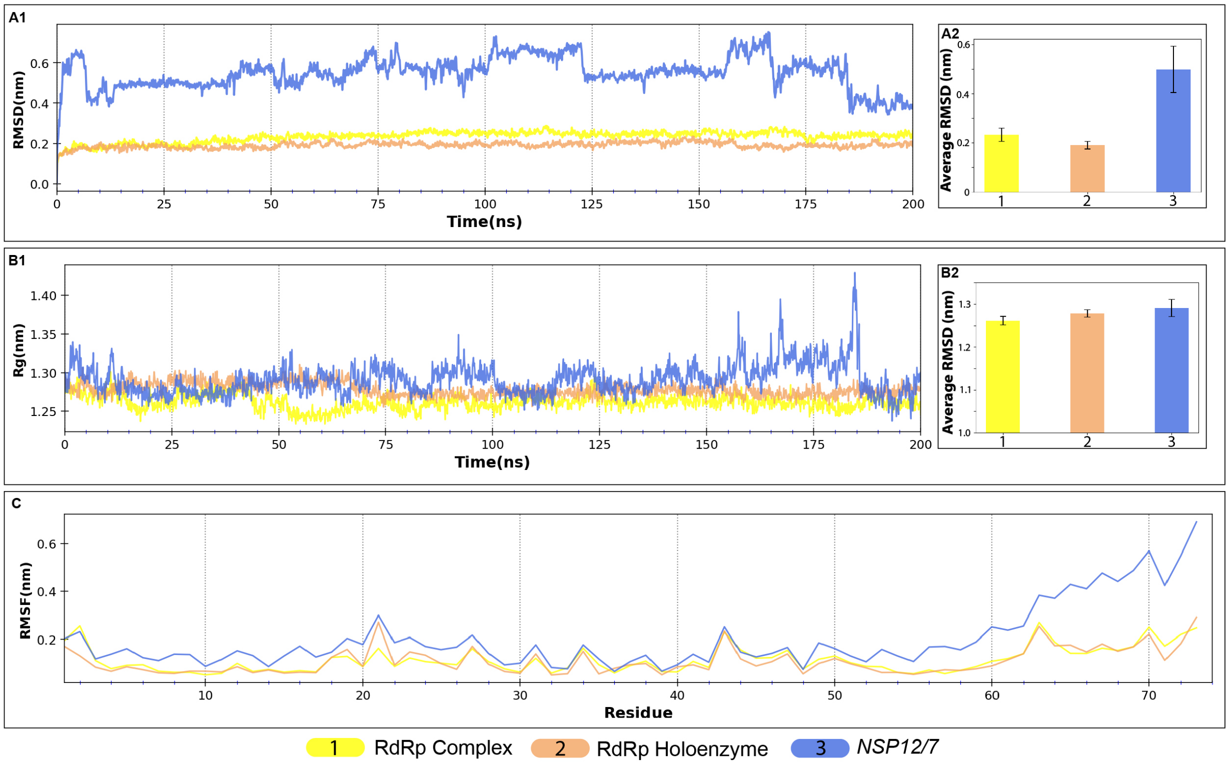


**Figure S5** showing the MD simulations analyses for *NSP12* in *NSP12/7* complex. **A1.** RMSD, **B1.** Rg, **C.** RMSF. The bar charts **A2** and **B2** placed adjacent to **A1** and **B1** illustrate the statistical representation of the data.

**Table S1:** ***In vitro*** **biochemical assays to delineate the roles of** ***NSP12/8/7*** **during the formation of the replication complex**

| **Experimental Set** | **Template-Primer Complex** | ***Experimental Conditions*** | |
|---|---|---|---|
| **Set # 1** | RNA-$U_{10}$ + RNA-P* for 60’ | *None* | *None* |
| | RNA-$U_{10}$ + RNA-P* | *NSP12 for 60’* | *None* |
| | RNA-$U_{10}$ + RNA-P* | *NSP12/NSP8/NSP7 for 60’* | *None* |
| | RNA-$U_{10}$ + RNA-P* | *NSP12 for 15’* | *NSP7 for 45’* |
| | RNA-$U_{10}$ + RNA-P* | *NSP12 for 30’* | *NSP7 for 30’* |
| | RNA-$U_{10}$ + RNA-P* | *NSP12 for 15’* | *NSP8 for 45’* |
| | RNA-$U_{10}$ + RNA-P* | *NSP12 for 30’* | *NSP8 for 30’* |
| | RNA-$U_{10}$ + RNA-P* | *NSP12 for 15’* | *NSP7/NSP8 for 45’* |
| | RNA-$U_{10}$ + RNA-P* | *NSP12 for 30’* | *NSP7/NSP8 for 30’* |
| **Set # 2** | RNA-$LU_8$* for 60’ | *None* | *None* |
| | RNA-$LU_8$* | *NSP12 for 60’* | *None* |
| | RNA-$LU_8$* | *NSP12/NSP8/NSP7 for 60’* | *None* |
| | RNA-$LU_8$* | *NSP12 for 15’* | *NSP7 for 45’* |
| | RNA-$LU_8$* | *NSP12 for 30’* | *NSP7 for 30’* |
| | RNA-$LU_8$* | *NSP12 for 15’* | *NSP8 for 45’* |
| | RNA-$LU_8$* | *NSP12 for 30’* | *NSP8 for 30’* |
| | RNA-$LU_8$* | *NSP12 for 15’* | *NSP7/NSP8 for 45’* |

| | | | |
|---|---|---|---|
| | RNA-LU8* | *NSP12 for 30'* | *NSP7/NSP8 for 30'* |

**Table S2:** ***The list of RNA binding cavity residues***

| **Residues of *NSP12* selected for cavity calculation using ANA2** |
|---|
| Ile494, Asn496, Asn497, Lys500, Ser501, Gly503, Gln541, Asn543, Lys545, Ala547, Ile548, Ala550, Lys551, Arg553, Arg555, Val557, Val560, Arg569, Gln573, Leu576, Lys577, Ala580, Val588, Ile589, Gly590, Thr591, Ser592, Lys593, Phe594, Tyr595, Trp598, Met601, Thr680, Ser681, Ser682, Gly683, Asp684, Ala685, Thr686, Thr687, Ala688, Tyr689, Leu758, Asp761, Phe812, Cys813, Ser814, Gln815, Pro832, Arg836, Ile837, Ala840, Lys849, Arg858, Ser861, Leu862, Ile864, Asp865, Met924, Thr929 |

**Table S3:** ***The RNA binding tunnel volume for the 6 complexes***

| **Complex** | **Cavity Volume (Å$^3$)** |
|---|---|
| RdRp Complex with RNA | 7139 |
| *NSP12* with RNA | 8116 |
| RdRp holoenzyme | 7560 |
| *NSP12* | 7734 |
| *NSP12/8* | 6789 |
| *NSP12/7* | 6830 |
